# Identifying Networks within an fMRI Multivariate Searchlight Analysis

**DOI:** 10.1101/2025.04.21.649903

**Authors:** Medha Sharma, Marc N. Coutanche

**Author notes:** Corresponding author: Marc N. Coutanche, LRDC, Murdoch Building, 3420 Forbes Ave, Pittsburgh PA 15213.

## Abstract

There is great interest in understanding how different brain regions represent information across space and time. Information-based searchlight analyses systematically examine the information encoded within clusters of functional magnetic resonance imaging (fMRI) voxels across the brain. Significant searchlights contain information that can be used to decode conditions of interest, but significant discriminability can be achieved in a variety of ways. We have developed and report on a new analysis method that can identify sub-networks of searchlights. Notably, unlike methods that collapse trials by condition, such as Representational Similarity Analysis, our method groups searchlights based on them having similar temporal changes in information. We present this method and apply it to fMRI data collected as participants viewed words, faces, shapes, and numbers. After running a searchlight analysis with a 4-way Gaussian Naive Bayes (GNB) classifier, the accuracy vector was submitted to a multi-subject Independent Component Analysis (ICA) to group searchlights based on their decoding timeseries. The ICA identified seven components (sub-networks) of searchlights. These networks identified sets of brain areas that have been commonly associated with the processing of faces, words, shapes and numbers.

For instance, two of the components drew strongly on the face-processing network, including fusiform cortex. Switching the classification scheme to ‘faces’ versus ‘non-faces’ reconfigured the observed network to reflected face-related systems. These results demonstrate that this method can divide searchlight maps into meaningful components.

## Identifying Networks within an fMRI Multivariate Searchlight Analysis

Information in the brain is reflected in different formats, including differing activation levels (Friston et al., 1994, 1995), subtle distributed patterns of activity (Haxby et al., 2001), unique connectivity footprints (Finn et al., 2015), and more. One functional magnetic resonance imaging (fMRI) analysis method for interrogating distributed activity patterns is multivariate pattern analysis (MVPA). MVPA involves the use of machine learning classifiers and related techniques to discriminate patterns of activity associated with different conditions. This can often identify information that is not apparent from univariate analyses that rely on the general linear model (Coutanche, 2013). Regions across the brain often differ in how a given set of information is represented.

A common application of MVPA is an ‘information brain mapping’, or ‘searchlight’, approach that sequentially analyzes clusters of voxels surrounding each searchlight center (Kriegeskorte et al., 2006). In its most common form, after running a searchlight analysis, the decoding performance (i.e., classifier accuracy) for each searchlight is allocated to its central voxel, generating a map of decoding accuracies across the brain. This is a useful way to identify regions that successfully discriminate conditions of interest (e.g., 70% accuracy where chance might be 50%). Although it is useful to identify the presence of discriminable patterns, one limitation of a single classification value is that different regions can have the same value (e.g., 70%) for distinct reasons. For instance, it might be possible to discriminate objects based on color or shape (Coutanche & Thompson-Schill, 2015) based on vastly different information across brain areas.

Representational similarity analysis (RSA) provides one solution to this concern by assessing the similarity between activity patterns that belong to different stimuli or conditions, to help reveal the basis for any discrimination. For instance, in the example above, green spheres and orange spheres will have very similar patterns in an exclusively shape-sensitive region, but not in an exclusively color-sensitive region (and *vice versa* for a green sphere and green cylinder). One disadvantage of RSA is that it eliminates the temporal dimension by collapsing the timeseries, which removes inter-trial differences within each condition. Such differences can reflect relevant differences between items (e.g., bright versus dull orange spheres) and fluctuating cognitive states (e.g., more versus less attention; Aly & Turk-Browne, 2016).

Collapsing the timeseries for each condition can improve the signal-to-noise ratio for certain questions but in the process eliminates important information in across-time variation (Coutanche, 2024). RSA’s aggregation of information into a single measure can also remove information from multiple dimensions (Davis & Poldrack, 2013). The method introduced here avoids these concerns by applying independent component analysis (ICA) directly to the decoding accuracy timeseries of searchlights, without collapsing values across time.

We introduce a new method that identifies brain-wide sub-networks of searchlights based multivariate pattern information across time. We apply the method to data collected as participants viewed images from four visual categories and report the resulting identified networks as proof of principle.

## Methods

### Subjects

The analyzed data were collected from twenty subjects from the Pittsburgh community (12 females, mean (M) age =22.5, standard deviation (SD)=3.9). All participants were right-handed, native English speakers without any neurological, psychiatric, learning or attention disorders. Informed consent was obtained from all participants, and the study was approved by the University of Pittsburgh Institutional Review Board. The data were collected as part of a previous investigation into number processing (Koch et al., 2023).

### Image acquisition

Participants were scanned using a Siemens fMRI 3T head-only Allegra magnet with a 32-head coil which included a mirror device for presentation of fMRI stimuli. The first session was T-1 weighted anatomical scan (TR = 1540 ms, TE = 3.04 ms, voxel size =1.00 x 1.00 x 1.00 mm, 192 slices). T-2 weighted functional scans were then collected, with slices collected in an interleaved, ascending order, with no skips between slices (TR = 2000 msec, echo time = 25 msec, flip angle = 70°, isotropic voxel size = 3.125 × 3.125 × 3.125 mm, 36 slices, in-plane resolution = 64 × 64, field of view = 200 × 200 mm). The functional scans were collected in four runs with 80 volumes each.

### fMRI preprocessing

Analysis of Functional Imaging (AFNI; Cox, 1996) was used to pre-process functional images with slice timing correction, motion correction, registration, high pass filtering and scaling voxel values. The functional data were standardized into MNI space using the MNI-152 T1 template. The Princeton MVPA toolbox (Detre et al., 2006) was used to import the data into MATLAB for subsequent analyses.

### Experimental task

While in the fMRI scanner, participants completed a match vs non-match judgement task (Emerson & Cantlon, 2015) with four stimuli conditions – words, faces, shapes, numbers (quantities & numerals). Each run contained two blocks for each of the four conditions. The blocks contained three trials (two seconds), each separated by two seconds of fixation. Blocks were separated by an eight-second fixation period and were presented in a pseudo-random order so that a block of every condition was presented before proceeding to the second set. During the words and faces conditions, words and faces were presented on either side of the fixation cross in lower/upper case and different orientation, respectively. Participants were asked to determine if the stimuli were the same or different. For the shape condition, participants judged if shapes (geometric and outlines of objects) were the same items. For the number condition, participants indicated whether the Arabic numeral represented the same number as dots in an array.

### Searchlight analysis

A roaming searchlight (3-voxel radius) was applied across the brain. The voxels in each searchlight were used as features in a Naive Bayes classifier for 4-way classification of the timepoints (TRs), based on the block label – words, faces, shapes, numbers. The four runs of the functional data were used as a 4-fold cross-validation set. The accuracies from the four folds were averaged and assigned to the center voxel of the searchlight. A map of significant searchlights was identified with a t-test applied to the twenty participants’ standardized accuracy maps compared to chance (0.25). Because this procedure identifies searchlights for the subsequent step, which itself includes a thresholding step, we employed a liberal p-value of 0.05 (uncorrected) in selecting the searchlights. The significant searchlights were advanced to the next step, which analyzed their timeseries of binary classification accuracies.

### Identifying sub-networks

The above analysis identified 15,544 significant searchlights, giving an accuracy matrix of 15,544 x 160 timepoints for each subject. We first concatenated the time points of all the participants to give a 15,544 voxel x 3,200 timepoint matrix for an ICA (Hyvärinen & Oja, 2000), in order to identify different components/clusters at the group level. To select the final number of components, a Principal Component Analysis (PCA) was used to visualize the variance-explained through a Scree plot. The presence of an elbow where the variance started decreasing less was used to identify the appropriate number of components (Calhoun et al., 2009).

Of the searchlights selected up to this point, a subset were advanced based on their relationship with at least one component: searchlights were retained if their component weight was greater than two standard deviations from the mean component weight (Calhoun et al., 2001; Kairov et al., 2017). In order to further improve robustness, we removed clusters with fewer than 10 voxels (searchlights) for each component.

### Component generalizability

To verify that any identified components are generalizable, we divided the participants into two groups and ran ICA for each group. We compared the percentage overlap in voxels being present in the same component across the samples (2 x common voxels / total voxels in both samples). We repeated this process 10 times, each with a new random division into two groups. To calculate significance, a null distribution of overlap values was created by randomizing the timeseries and running the ICA and overlap analyses. This was performed 100 times, to generate a null distribution of the probability for a given voxel to fall within the same component.

## Results

### Identification of searchlights

To first identify discriminable searchlights, we ran a searchlight procedure to classify the four visual categories in each participant’s brain. The classification accuracy of each searchlight was tested against chance (0.25) through a t-test at the group level (Figure 1). As described above, a liberal p < 0.05 uncorrected threshold was employed to inclusively select relevant searchlights prior to the next analysis step and additional thresholding.

**Figure 1:**
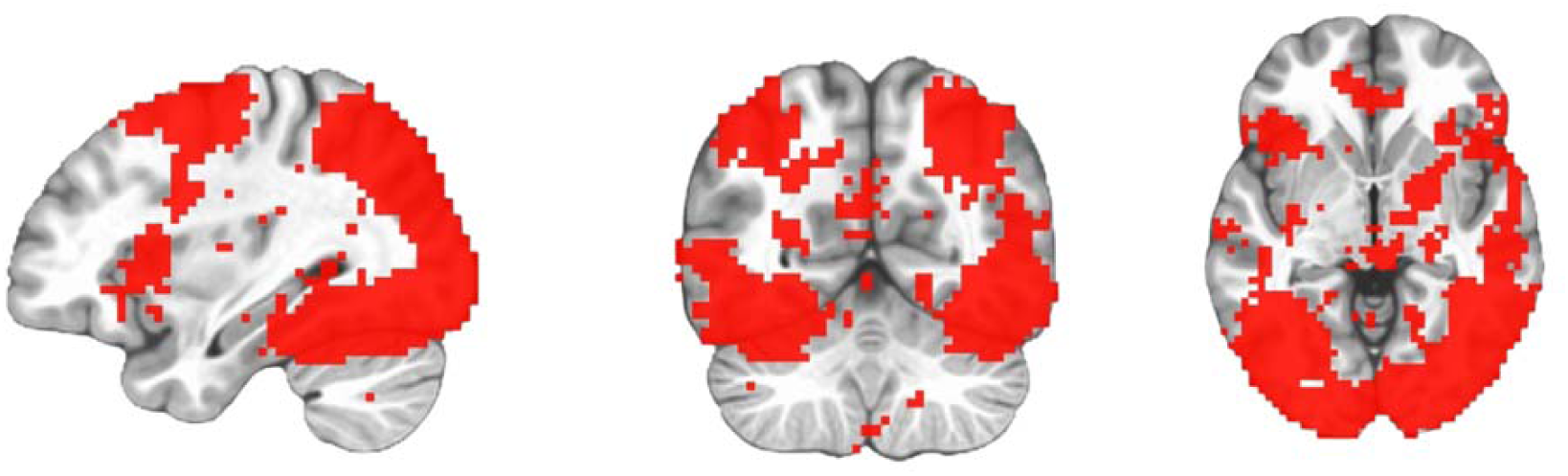
T-test results of the group searchlight accuracies against 0.25. Slice coordinates: x=33 mm, y=-58 mm, z=-4 mm

### Identification of sub-networks

For each searchlight identified in the previous analysis, the binary accuracy vectors (length equal to the timeseries) from the participants were concatenated to produce an accuracy vector for the group. The resulting matrix served as input to ICA. A scree plot was used to determine the number of components (Figure 2). Seven components were identified based on the elbow reflecting the variance explained. Figure 3 shows the seven independent components identified (voxel counts: 614, 1016, 881, 1125, 1067, 858, 978).

**Figure 2:**
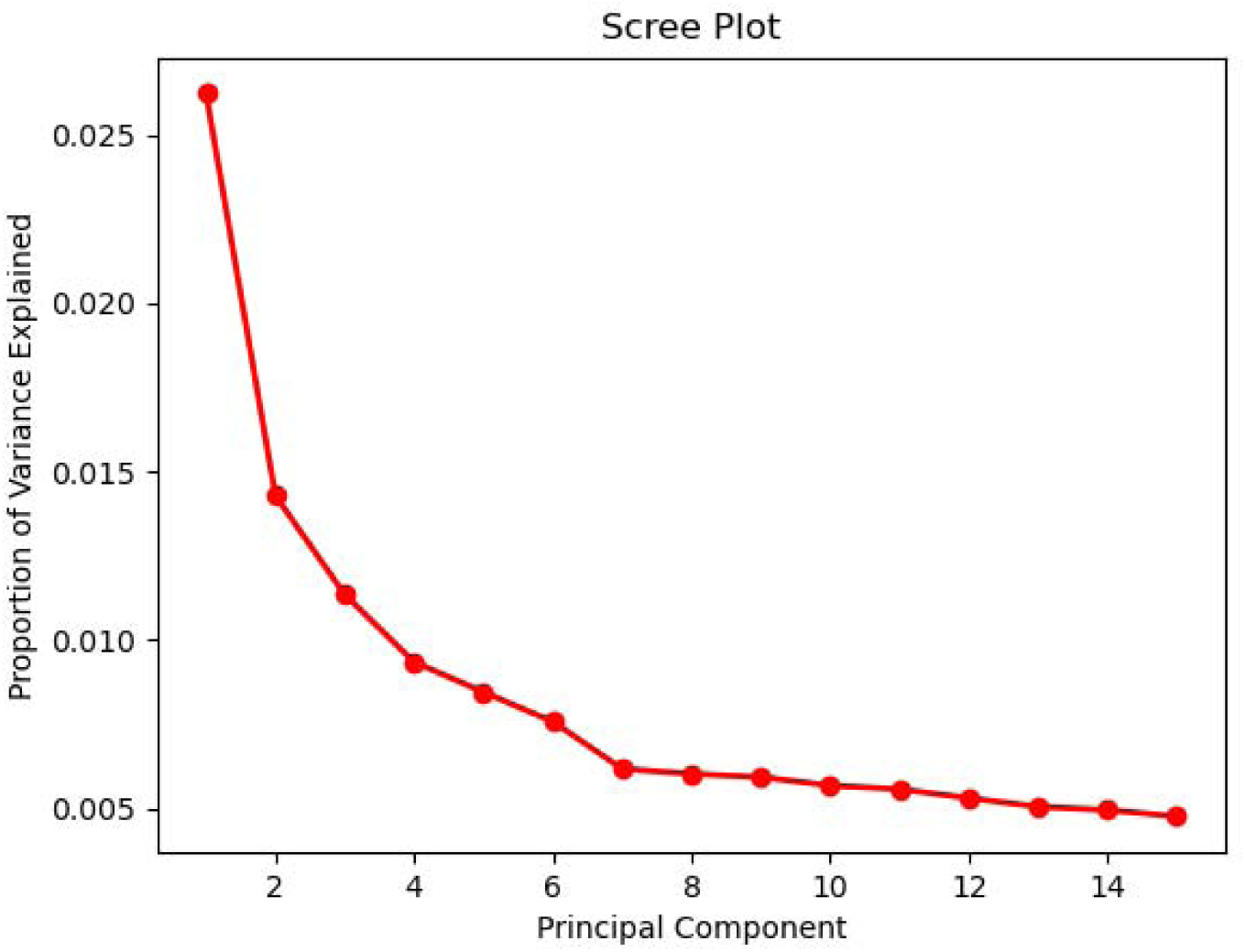
Scree Plot from Principal Component Analysis to determine number of components. The x-axis is capped at 15 for ease of visualization.

**Figure 3:**
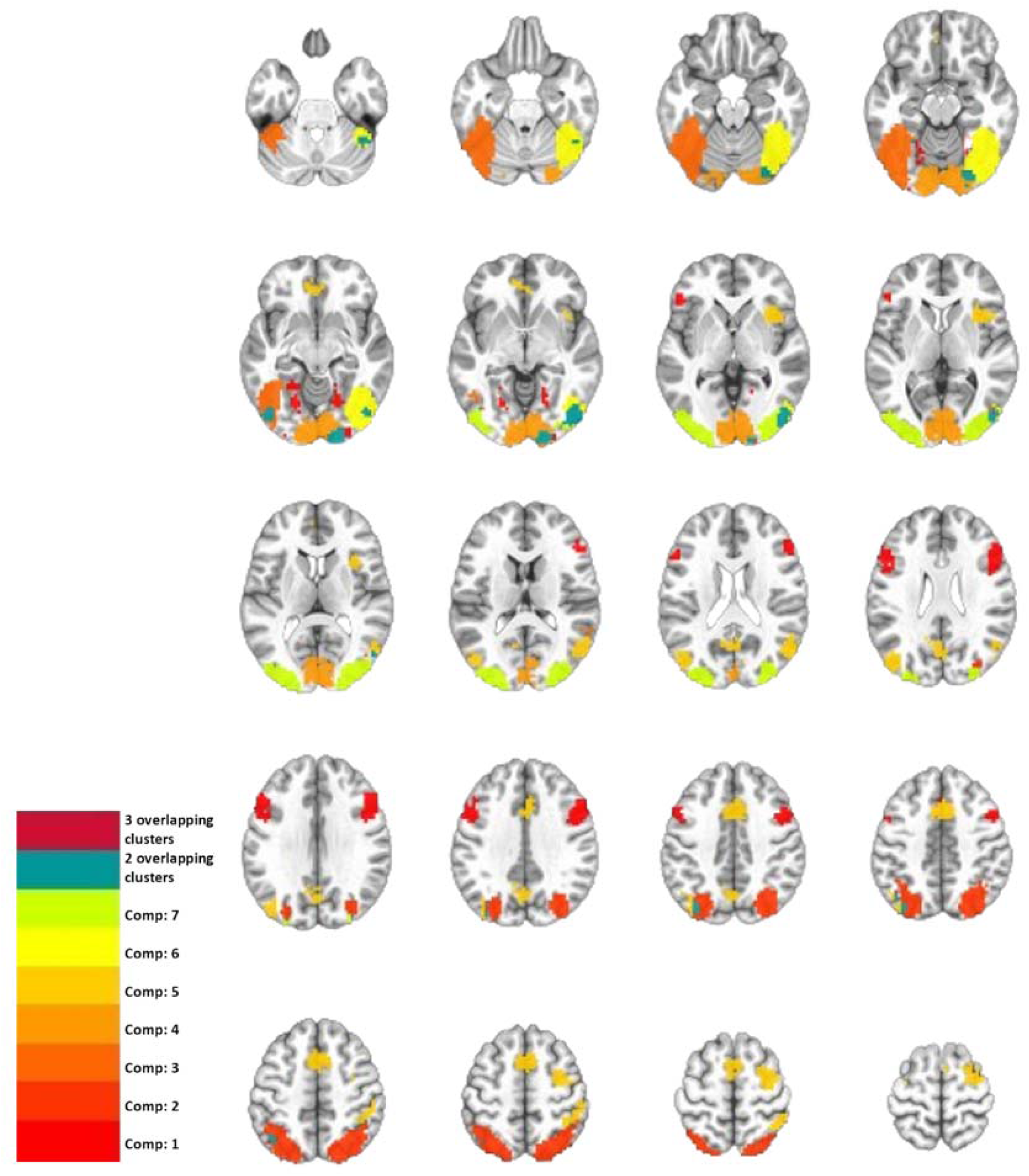
Montage of the seven Independent Components. This image shows the center voxels of the searchlights in the independent components. Axial slices go from z =-30mm with an increment of 5 mm up to z = 65mm.

### Individual Components

To examine the function of the searchlights in each component, we first examined the classification accuracies for each of the four conditions (Table 1; confusion matrices in Supplementary Materials).

**Table 1:** Average classification accuracy of each label. The accuracy value is averaged across the searchlights in each component. Chance performance is 0.25.

|  | Word | Face | Shapes | Number |
| --- | --- | --- | --- | --- |
| Component 1: | 0.2714 | 0.3140 | 0.3084 | 0.2878 |
| Component 2: | 0.2555 | 0.2745 | 0.2752 | 0.4077 |
| Component 3: | 0.2957 | 0.3913 | 0.3079 | 0.2818 |
| Component 4: | 0.2990 | 0.3346 | 0.3005 | 0.2766 |
| Component 5: | 0.2548 | 0.2912 | 0.2652 | 0.3284 |
| Component 6: | 0.3296 | 0.4808 | 0.2919 | 0.2716 |
| Component 7: | 0.3501 | 0.3675 | 0.2972 | 0.2761 |

Next, each component is described at the cluster level, with number of searchlights present, average accuracies by label, anatomical location, and associations in the literature extracted using Neurosynth (Yarkoni et al., 2011). Confusion matrices for each cluster are presented in the supplementary materials.

The clusters associated with each component are shown in Figures 4-10.

**Figure 4:**
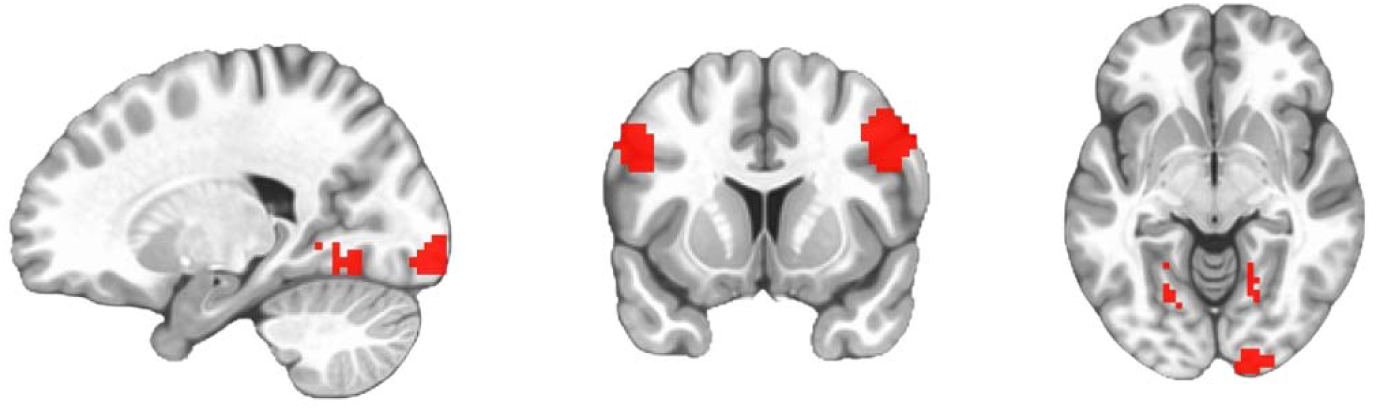
Component 1. Slice coordinates: x= 20 mm, y= 11 mm, z=-7 mm.

**Figure 5:**
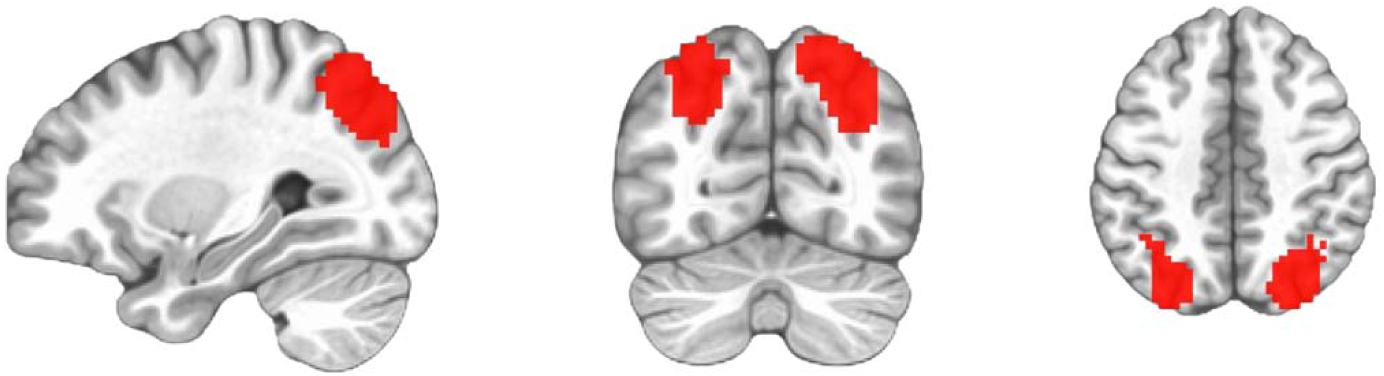
Component 2. Slice coordinates: x= 29 mm, y=-70 mm, z= 45 mm.

**Figure 6:**
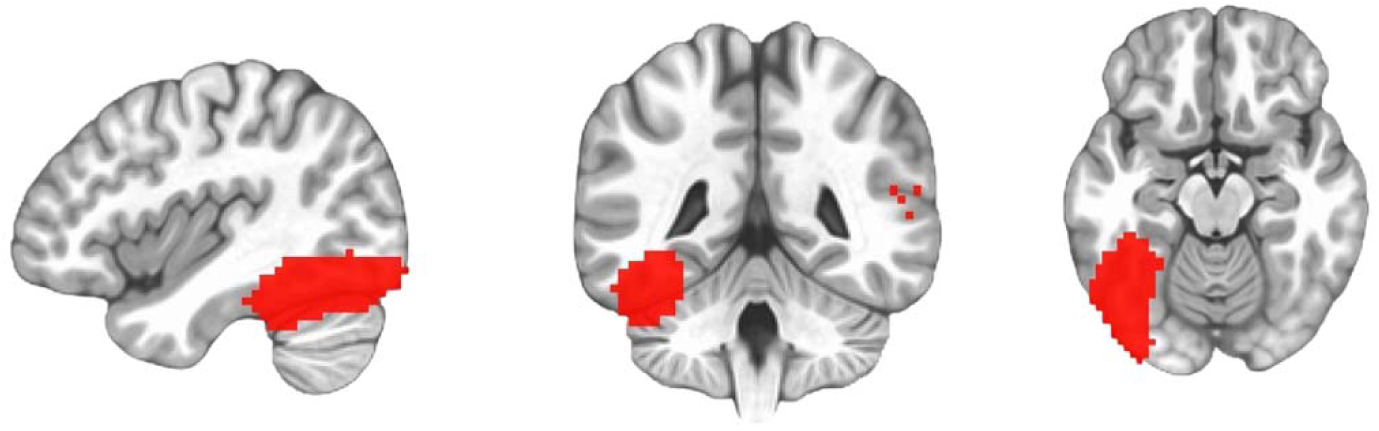
Component 3. Slice coordinates: x=-41 mm, y=-45 mm, z=-16 mm.

**Figure 7:**
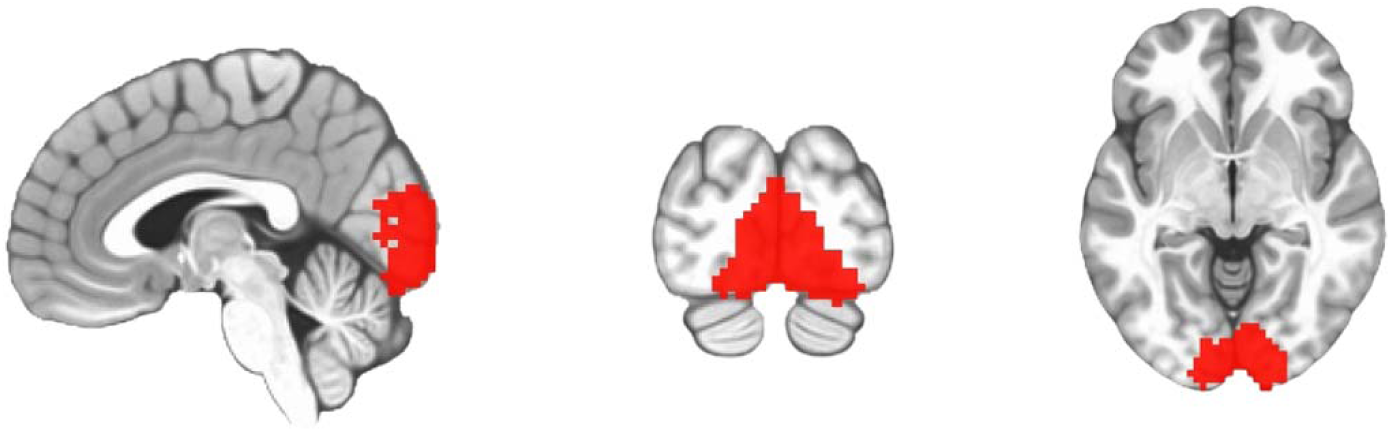
Component 4. Slice coordinates: x= 3 mm, y=-89 mm, z=-5 mm.

**Figure 8:**
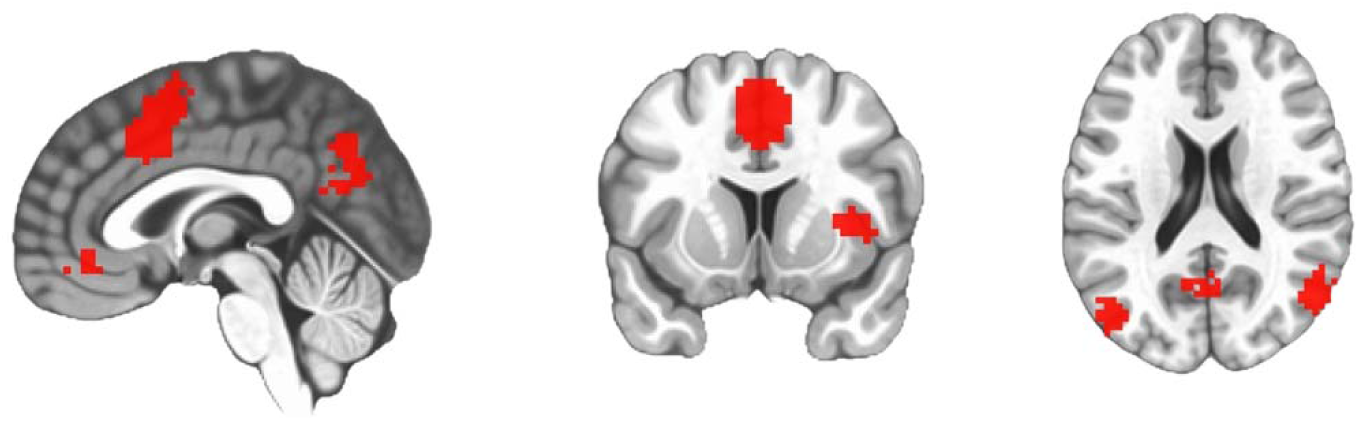
Component 5. Slice coordinates: x= 1 mm, y= 11 mm, z= 20 mm.

**Figure 9:**
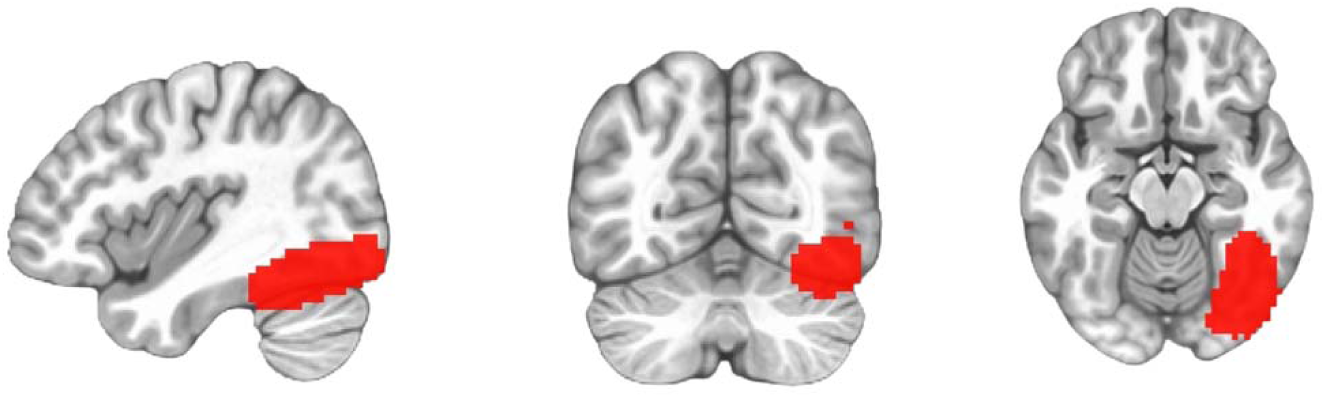
Component 6. Slice coordinates: x= 46 mm, y=-65 mm, z=-15 mm.

**Figure 10:**
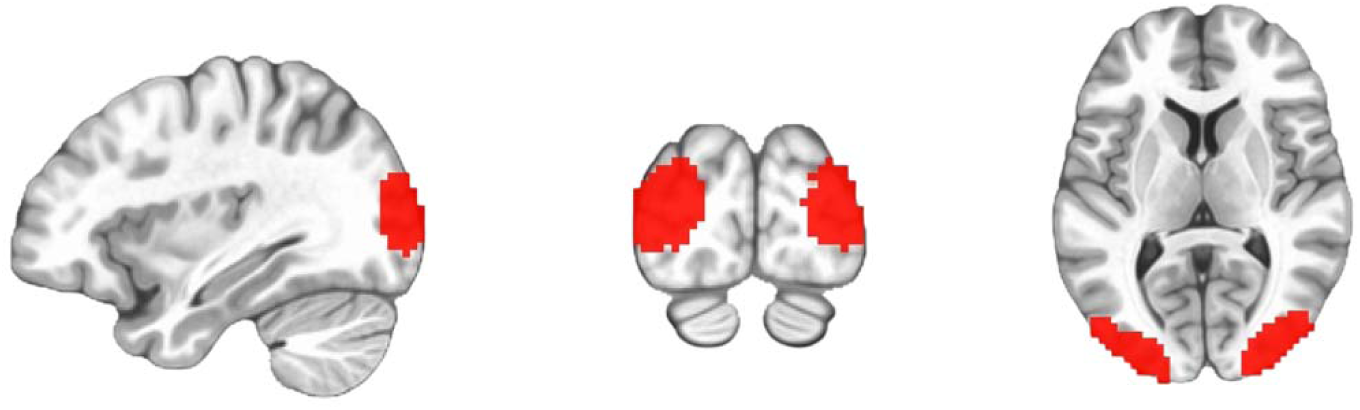
Component 7. Slice coordinates: x=-34 mm, y=-90 mm, z= 8 mm.

### Generalizability

To test whether the assignments of searchlights to components is robust and generalizable, we conducted the analysis within subsets of participants and asked if the searchlights were grouped together similarly across two groups of randomly selected participants. In every pair of subsamples (i.e., different ways of dividing the sample), searchlights were significantly more likely to be allocated to the same component, than would be expected by chance (all p<0.01), indicating that the assignment of searchlights to the same component is reliable across different groups of participants.

### Impact of classification scheme

To examine how the classification scheme impacts the resulting networks (for the same stimuli), we repeated the ICA procedure but with accuracy vectors from a ‘face versus non-face’ classification by labeling the words/shapes/numbers as the same ‘non-face’ condition. The resulting networks reconfigured to reflect brain systems underlying face processing, social cognition and emotional processing (Figure 11; Table 3). To statistically confirm this, we compared the number of ‘face’ (or ‘faces’) labels generated by the Neurosynth meta-analysis after the 4-way classification (11 ‘face’; 210 non-face labels) and after the face vs. non-face classification (19 ‘face’; 84 non-face labels). A chi-square analysis confirmed that the ICA using the ‘face versus non-face’ accuracy vector produced clusters with a significantly greater proportion of face associations than did the ICA with the four-way accuracy vector (*X^2^* = 15.17, p =.0001). Significance remained after removing duplicate face labels from each cluster and comparing the number of network clusters with at least one ‘face(s)’ label in Neurosynth after the 4-way classification (5 clusters with ‘face’; 19 without ‘face’) and face classification (6 clusters with ‘face’; 5 without ‘face’). A chi-square analysis confirmed that the ICA using the face versus non-face accuracy vector produced a significantly greater proportion of clusters that are associated with face-related processing in the neuroimaging literature (*X^2^* = 3.98, p =.046).

**Figure 11:**
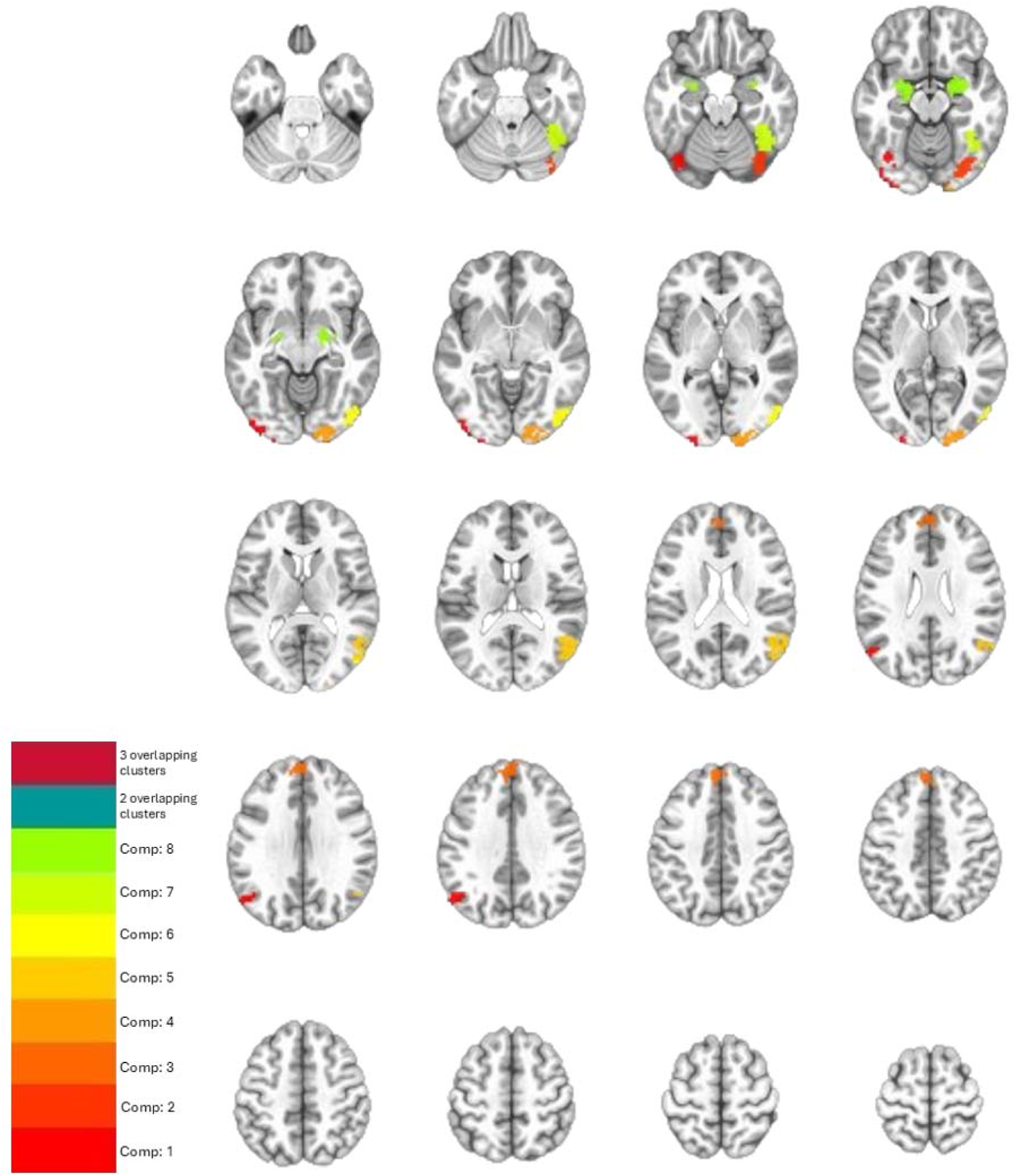
Montage of the eight independent components of searchlights generated from a face vs non-face classification. The voxels show the center of each searchlight. Axial slices: z=-30mm with an increment of 5mm up to z=65mm.

## Discussion

In this paper, we present and apply a new method for identifying sub-networks of information-based searchlights. The method uses ICA to group searchlights based on their decoding timeseries, identifying sub-networks from a searchlight analysis. We applied the method to data collected as participants undertook a discrimination task with four visual categories – faces, words, numbers, shapes. The method identified seven independent components with meta-analytical associations that were consistent with the examined conditions. Switching the basis for decoding to ‘faces versus non-faces’ shifted the components to reflect areas of the face network, supporting the method’s validity, accuracy and specificity.

A key challenge for any new method is demonstrating that its results are meaningful. In addition to the reconfiguration of networks when examining face decoding, repeatedly finding bilateral clusters suggests the method is correctly grouping searchlights with common neural representations. Specifically, components 1,2,4,5 and 7 of the 4-way classification included regions that are mirrored across the midline. The ICA and searchlight do not have information about which searchlights are hemispheric homologues, so the repeated appearance of laterality reflects the detection of common information time courses for certain bilateral regions. This provides validity to the results, as ICA is not just grouping nearby searchlights but is grouping distant regions with (some) commonalities in function.

We now briefly review the composition of each component. Due to the number of clusters presented, we focus on the largest clusters from each component and refer readers to the Results and Supplementary Materials for full details.

The first component featured clusters associated with across-condition processes, such as cognitive control, language, and reading words likely reflecting internal labelling of the stimuli and word condition (Chee et al., 2003). Notably, the third cluster is associated with ‘matching’, reflecting the discrimination task for the conditions. The remainder of the clusters were associated with general language, visual and memory processes. The second component was driven by number processing, which is relevant to one of the four conditions: numbers. Both clusters showed superior classification accuracy for the number condition, and associated terms were predominantly related to number processing and number-associated cognitive processing (e.g., calculation and phonology; Andin et al., 2015). The third component is related to word and face processing, with highest accuracy for face decoding, and associated terms being related to visual and word processing. This area is typically responsive during letter representation (Horie et al., 2012; Pernet et al., 2005), top-down processing of the words and pictures (Kherif et al., 2011) and face processing (Iidaka et al., 2006). The fourth component is one large cluster centered on early visual areas. It has similar classification accuracy for all four conditions and is associated with basic visual information across multiple categories (e.g., Coutanche et al., 2016). The fifth component is composed of clusters primarily in parietal, motor and prefrontal cortex, associated with working, motor control, and other processes. The sixth component has superior classification for faces and covers the fusiform gyrus, which is repeatedly reported in face studies (Haxby et al., 2001). The seventh component consists of three clusters located in visual areas with a preference for faces.

The presented method provides a new tool among several recent methods seeking to examine regions through the lens of information fluctuations over time (Anzellotti & Coutanche, 2018; Coutanche & Thompson-Schill, 2013, 2014; Huang et al., 2024). This method’s unique contribution is in the identification of sub-networks (i.e., grouping searchlights) rather than assessing informational connectivity from one seed region to other regions. The use of ICA, while powerful, also presents some limitations that are important to consider. For instance, it is necessary to determine the number of components that will be selected. We used the common approach of examining a data-driven scree plot of diminishing levels of variance-explained, but because there is no established boundary, the selection of the scree plot elbow can be subjective. Additionally, though it was clear for our design that we should include all subjects, studies with a more variable subject set (e.g., right-and left-handed, a wide age range), may wish to consider grouping participants by characteristics in order to identify robust components (which could then be compared). Another decision point is how to threshold the components weights when generating the spatial maps. This can be accomplished in various ways.

In conclusion, we have introduced and demonstrated an analysis method that identifies sub-networks of searchlights based on their shared fluctuations in discriminability. We hope this method will allow investigators across a range of fields to move beyond an initial decoding searchlight map and toward an understanding of the spatial organization of information across the brain.

## Data and Code Availability

The codebase for the analyses is available at https://github.com/Pitt-Cognim-Lab/IdentifyingNetworksUsingSearchlightMVPA. The data are available at https://openneuro.org/datasets/ds004594/versions/1.0.1.

## Author Contributions

M.S. – data curation, formal analysis, methodology, writing. M.N.C. – conceptualization, funding acquisition, methodology, writing, review & editing, supervision.

## Acknowledgments

The authors thank Julie Fiez and Melissa Libertus for their involvement in the design of the study generating the dataset.

## Supplementary Material

### Confusion matrices for ICA component clusters

The following confusion matrices show average predictions per class. Table 2 details the clusters.

**Table 2:**
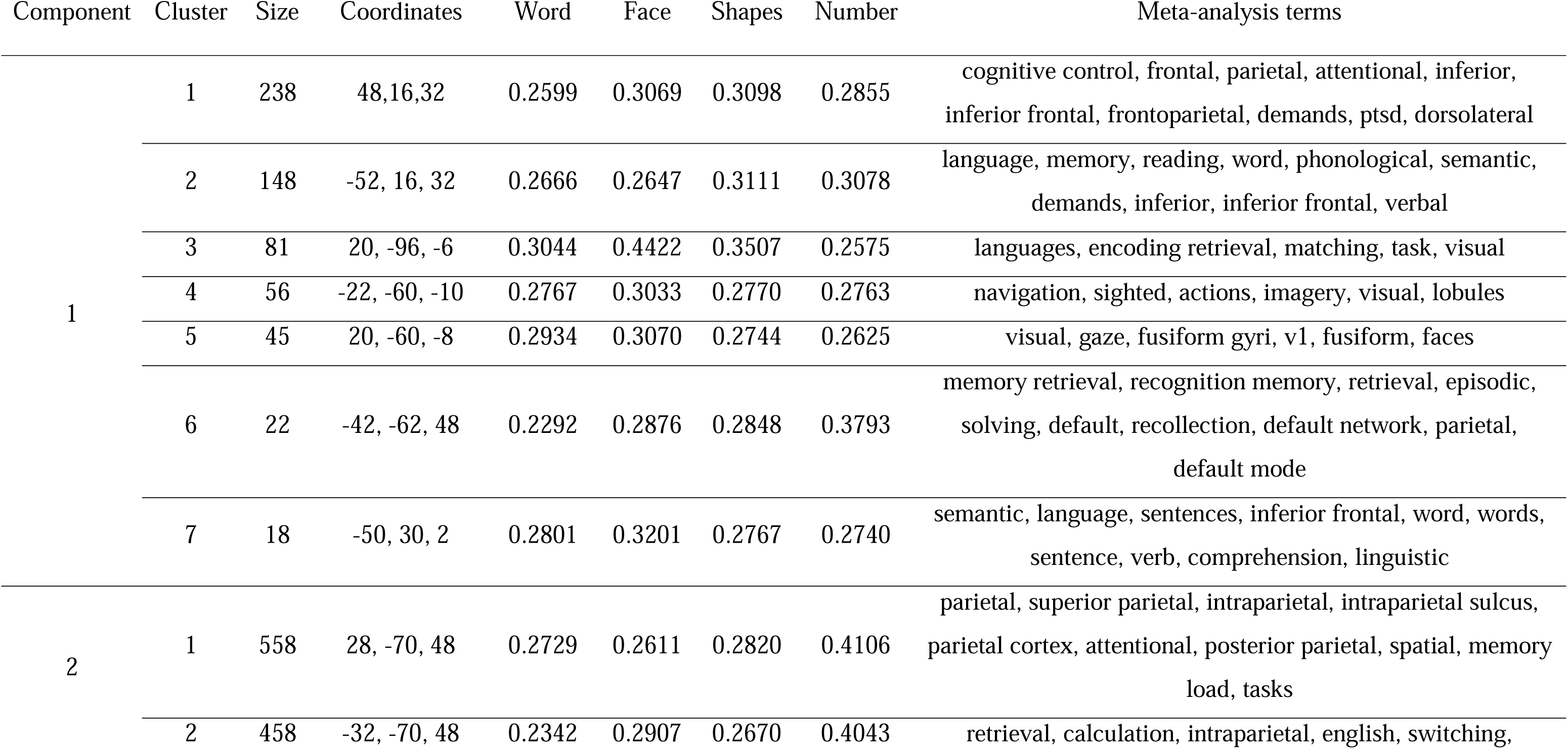
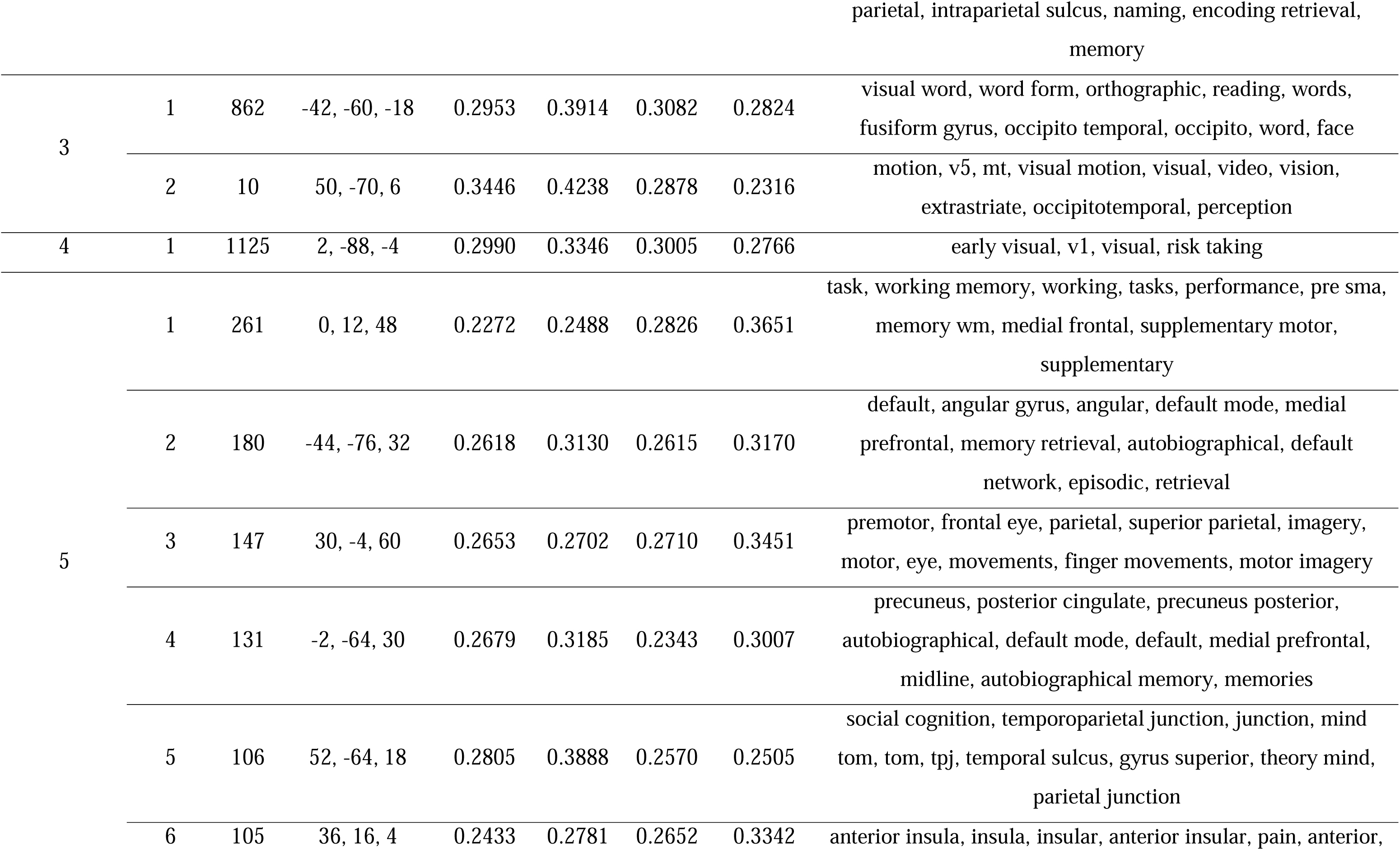
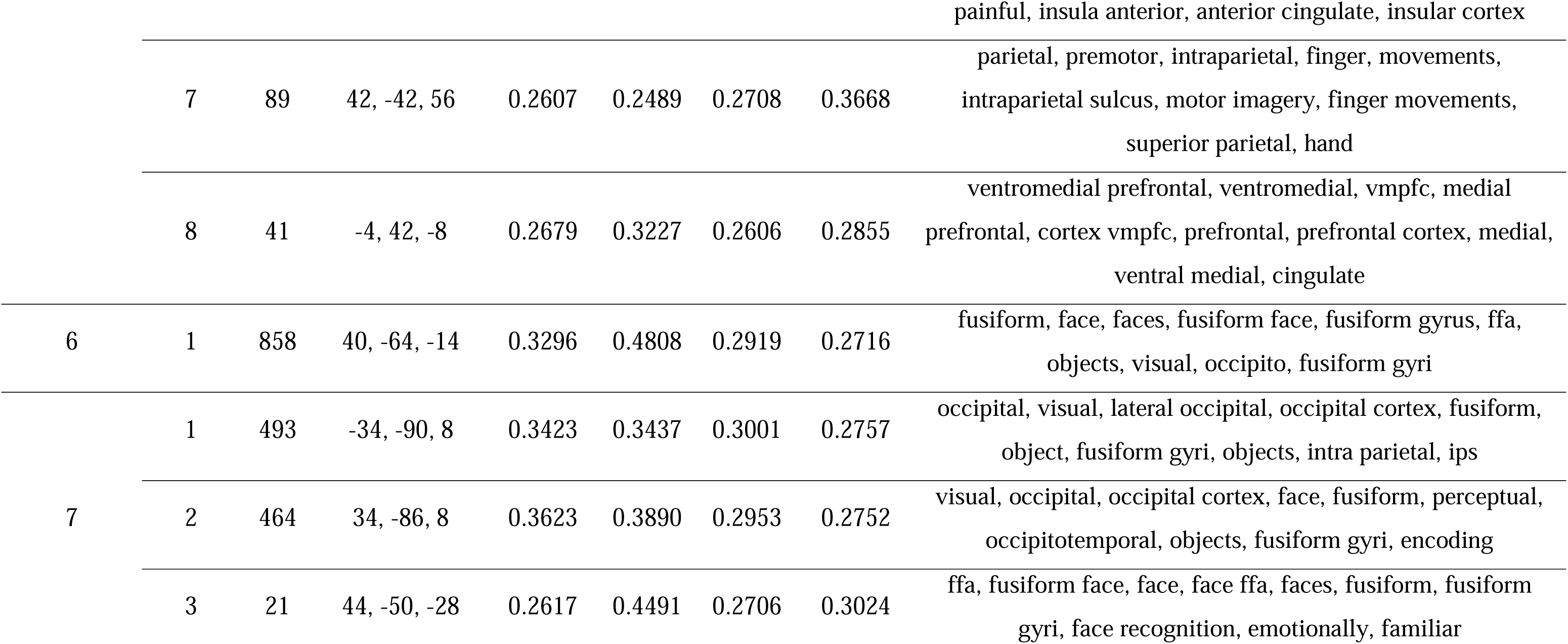
Clusters for the independent components from a four-way classification. The information includes cluster size, center of mass (MNI), class accuracy, and ordered terms from a Neurosynth meta-analysis. The top 10 terms are listed if the terms exceed 10.

**Table 3:**
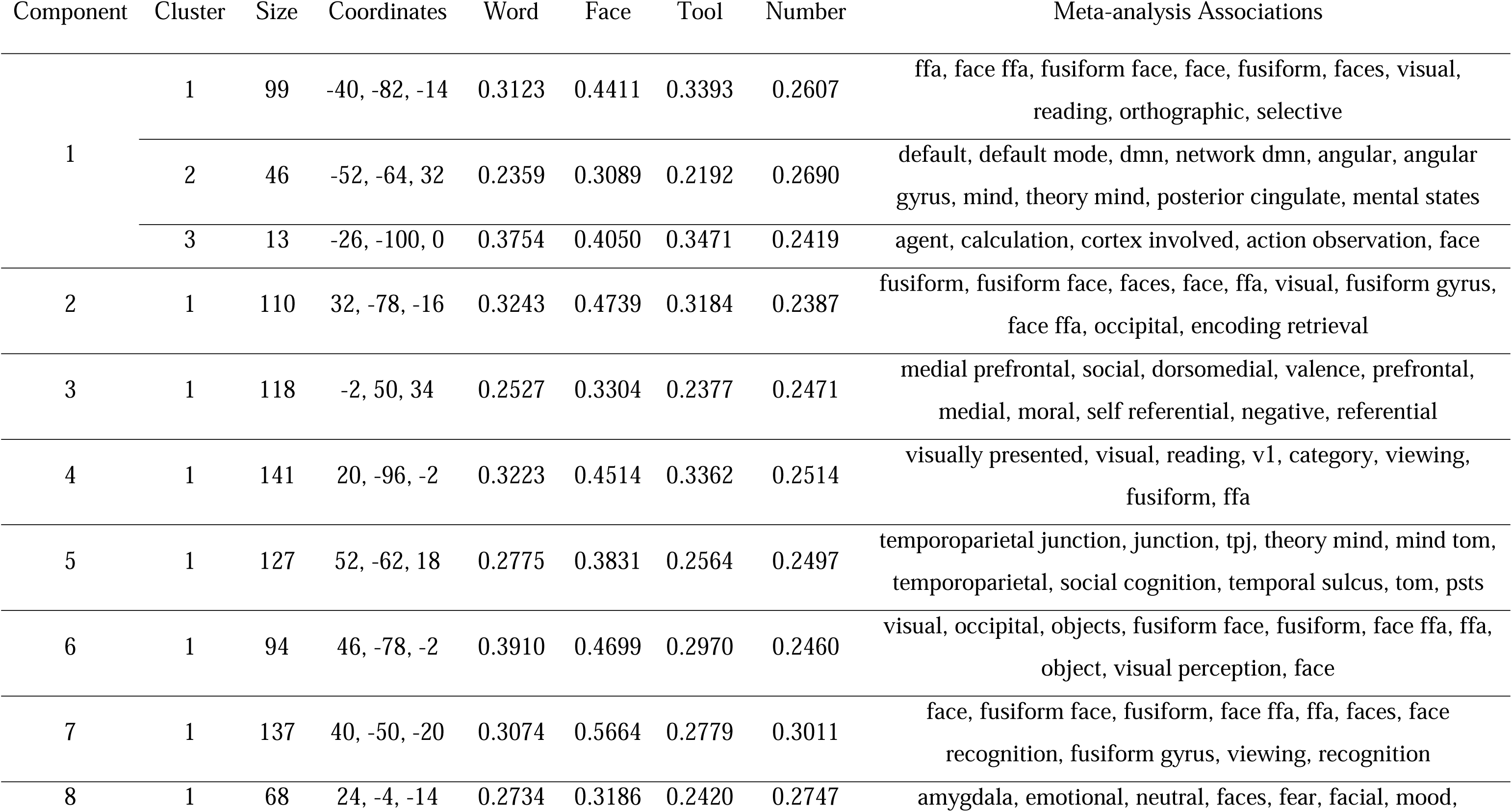
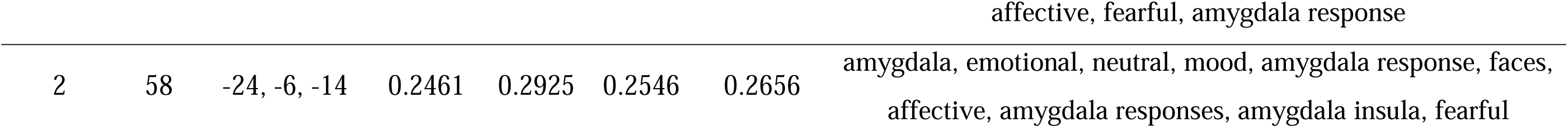
Clusters for the independent components from ‘face versus non-face’ classification. The information includes cluster size, center of mass (MNI), class accuracy, and ordered terms from a Neurosynth meta-analysis. The top 10 terms are listed if the terms exceed 10.

Component 1:

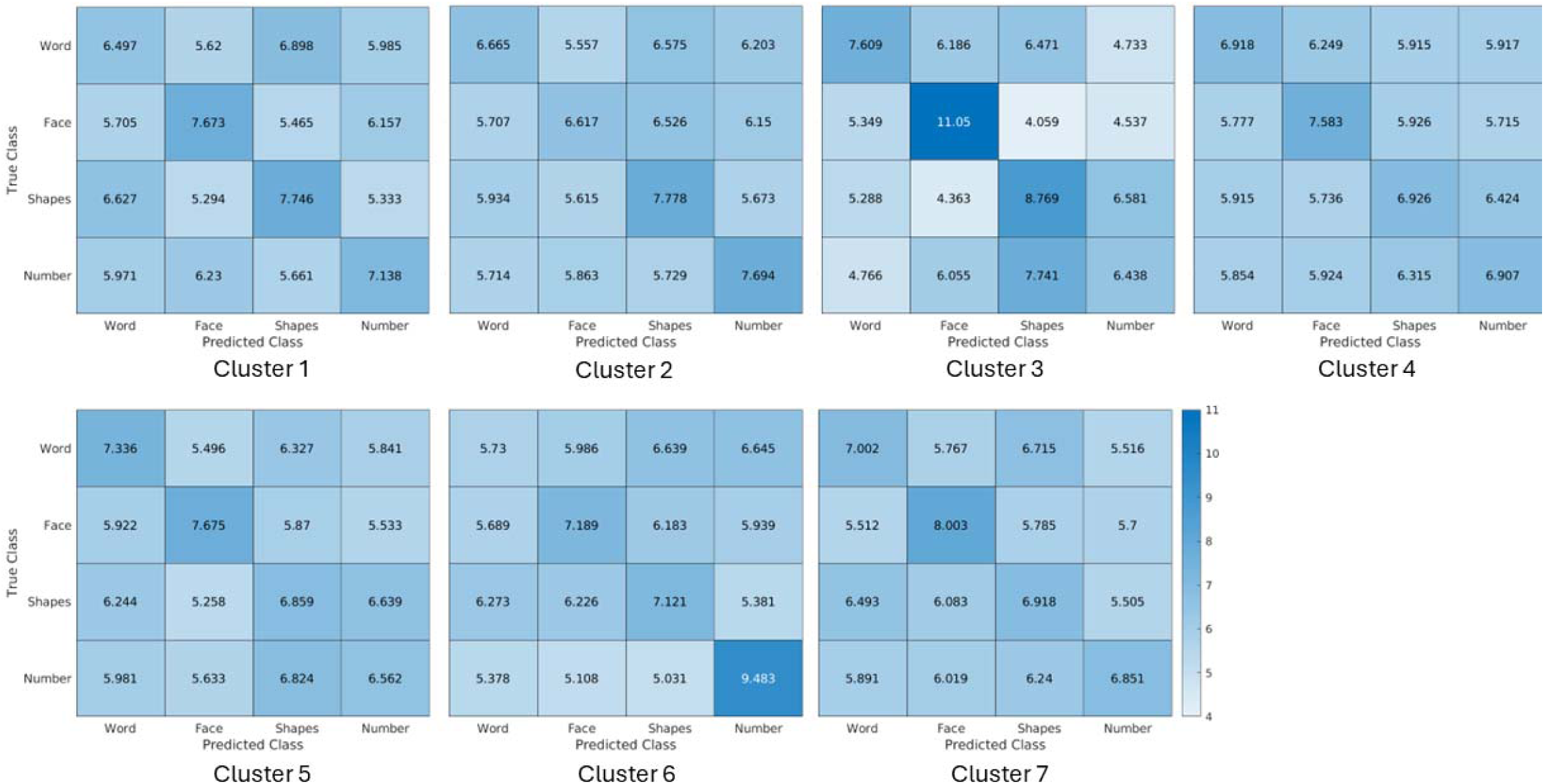

Component 2:

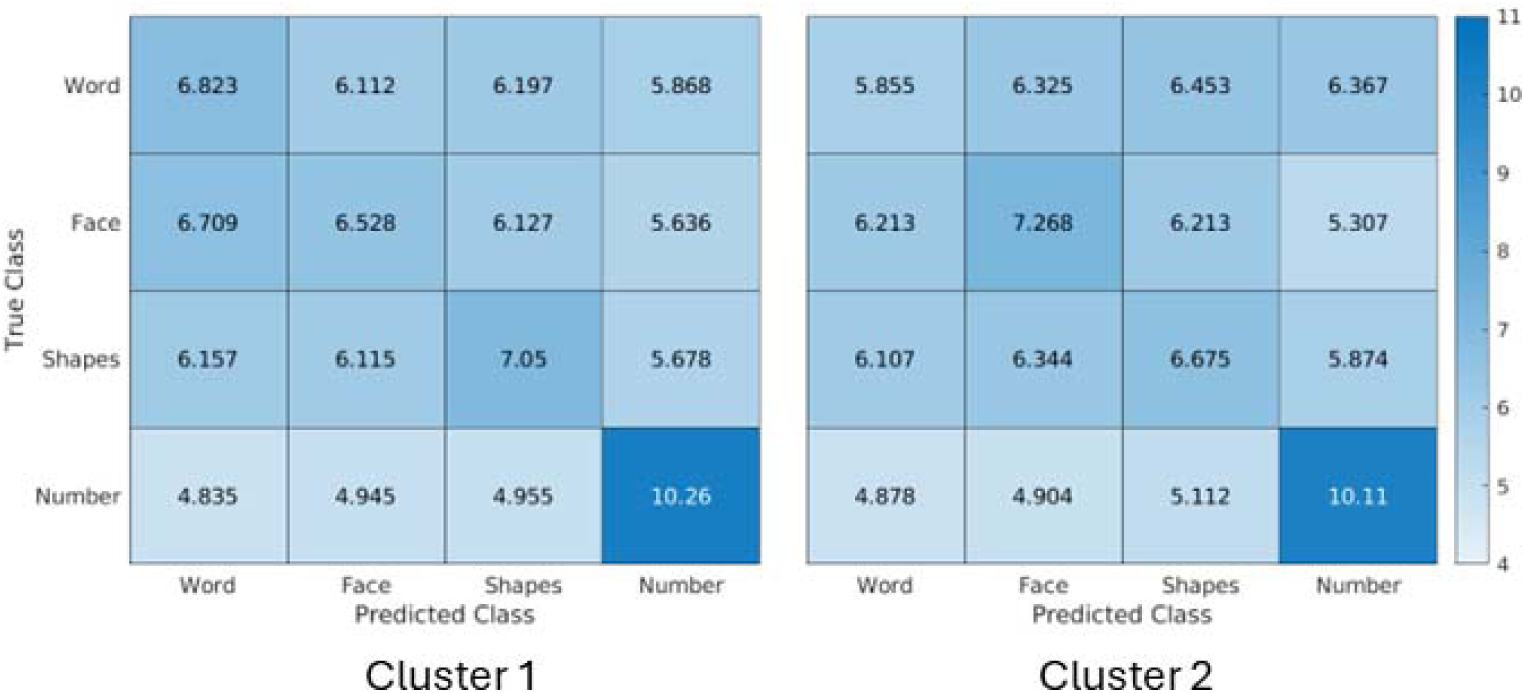

Component 3:

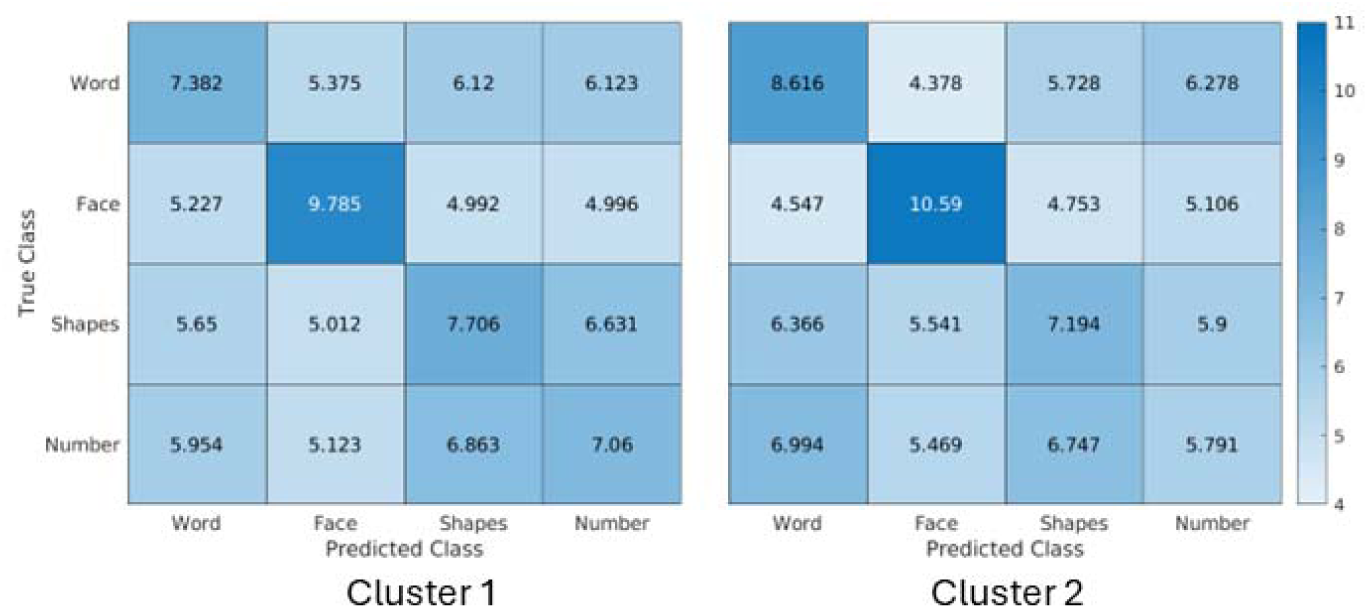

Component 4:

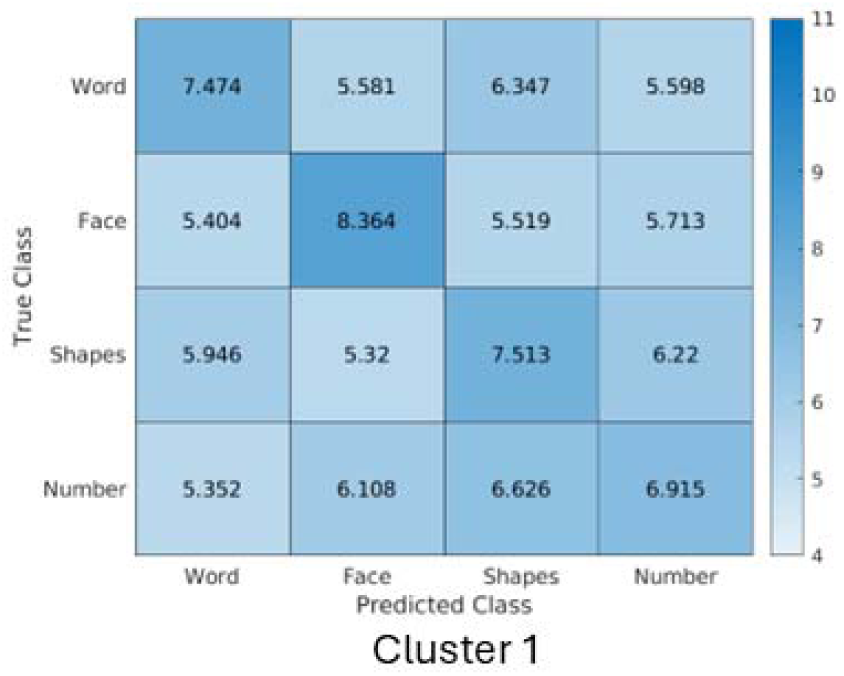

Component 5:

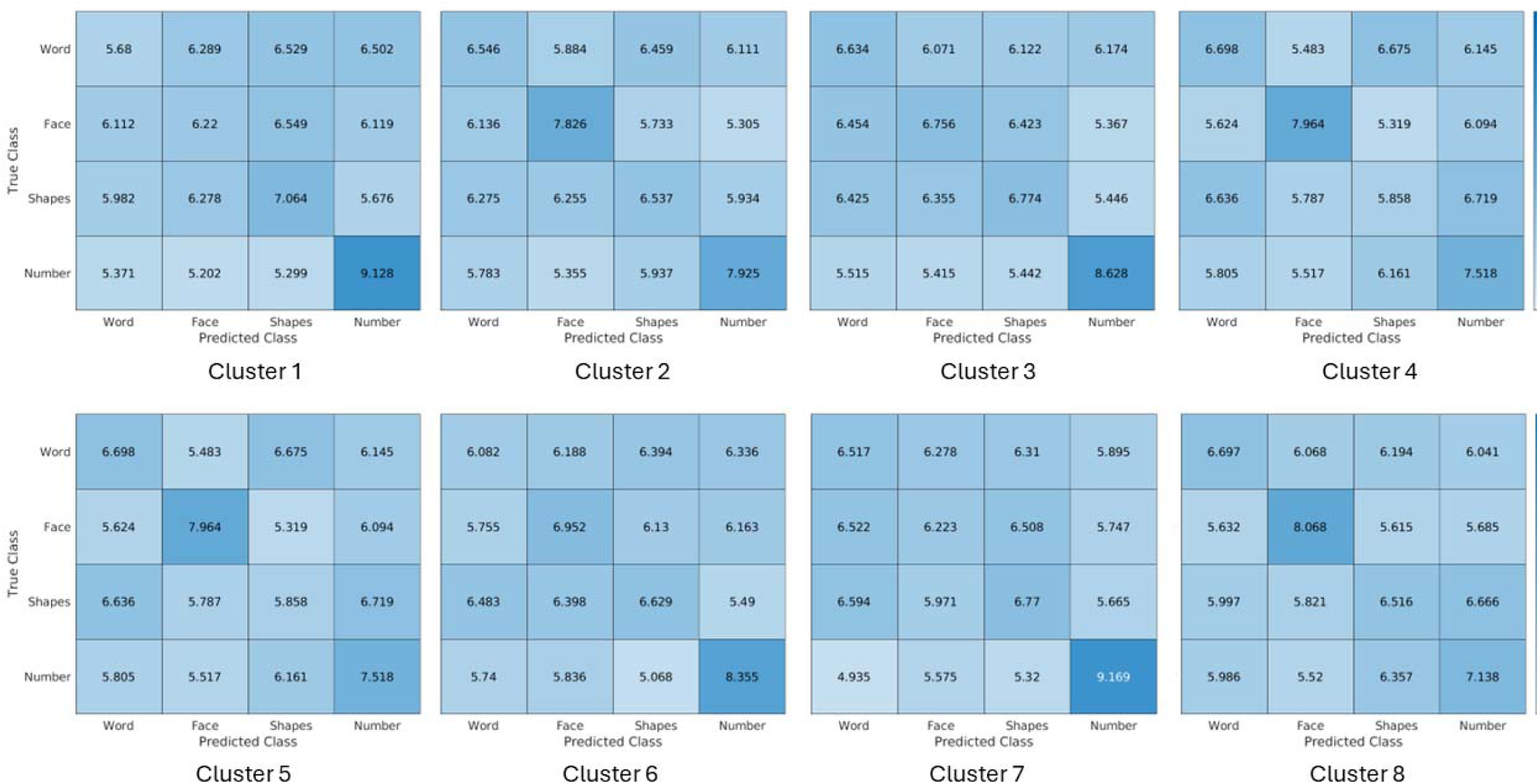

Component 6:

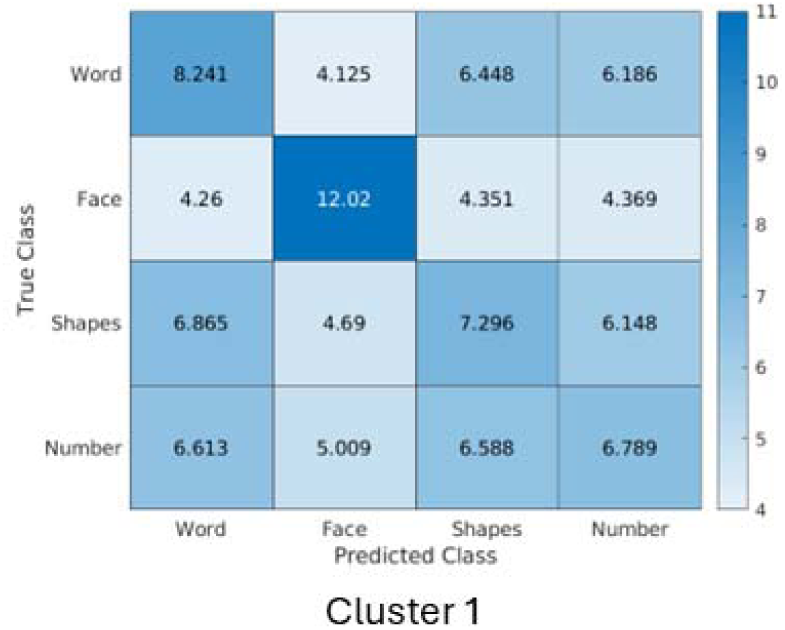

Component 7:

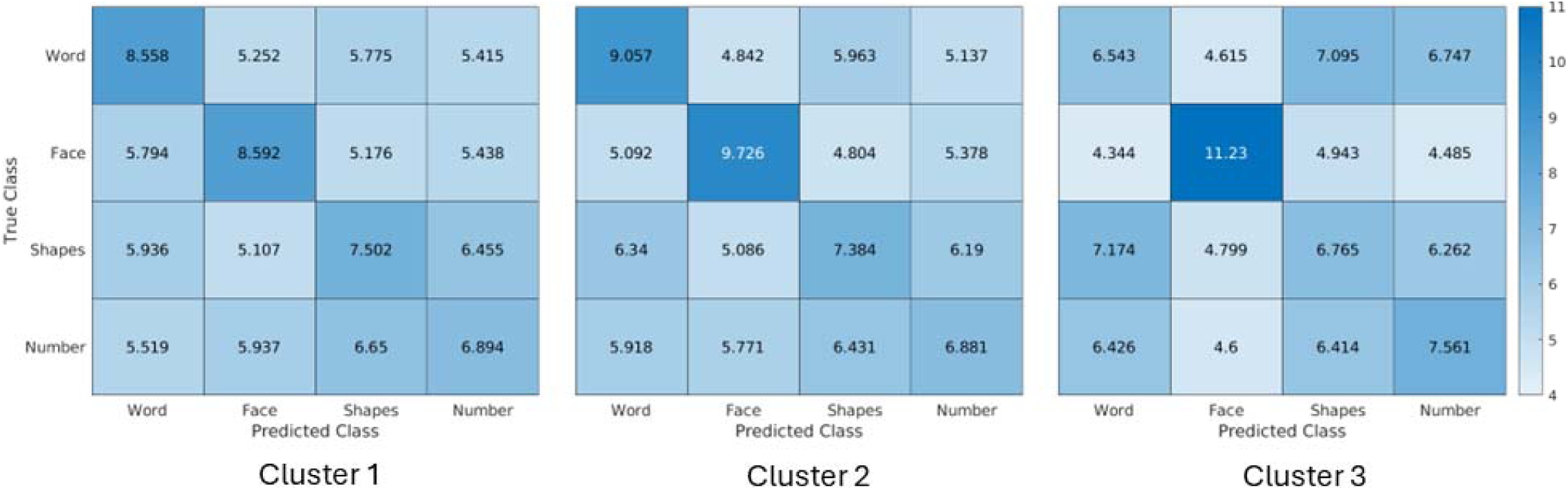

